# High-fidelity enhanced AsCas12a knock-in mice for efficient multiplexed gene editing, disease modeling and orthogonal immunogenetics

**DOI:** 10.1101/2024.03.14.585126

**Authors:** Kaiyuan Tang, Xiaoyu Zhou, Shao-Yu Fang, Erica Vandenbulcke, Andrew Du, Johanna Shen, Hanbing Cao, Jerry Zhou, Krista Chen, Shan Xin, Liqun Zhou, Shawn H. Lin, Medha Majety, Xingyu Ling, Stanley Z. Lam, Ryan Chow, Suxia Bai, Timothy Nottoli, Carmen Booth, Chen Liu, Matthew B. Dong, Sidi Chen

## Abstract

The advancement of CRISPR gene editing technology, especially the development of Cas9 knock-in mice, has significantly boosted the functional discovery of various genetics factors in diverse fields including genetics, genomics, immunology, and the biology of cancer. However, the pleiotropic effects on human disease and the complex nature of gene interaction networks require a knock-in mouse model capable of simultaneous multiplexed gene perturbation. Here, we present the generation and applications of Cre-dependent conditional and constitutive high-fidelity, enhanced AsCas12a (enAsCas12a-HF1) Rosa26-knock-in mice in the C57BL/6 background. With these mouse strains, we demonstrate highly efficient and multiplexed *in vivo* and *ex vivo* genome engineering as applied to lipid nanoparticle (LNP)-RNA-based liver protein targeting, AAV-based tumor modeling, and retrovirus-based immune cell engineering. By integrating with a dCas9-SPH CRISPR activation transgenic strain, we establish a simultaneous dual gene activation and knockout (DAKO) system that showcases the modular potential of these enAsCas12a-HF1 mice. Importantly, constitutive expression of enAsCas12a-HF1 does not lead to any discernable pathological differences as compared to the C57BL/6 background strain. These knock-in mice and the accompanying delivery methods would empower the deconvolution of complex gene interaction networks in broad areas of research.

## Introduction

The development of Clustered Regularly Interspaced Short Palindromic Repeats (CRISPR) technology has offered unprecedented capabilities in endogenous gene editing^1^. CRISPR technology utilized programmable RNA to direct endonuclease activity, which provides remarkable precision and simplicity to target any DNA or RNA sequences^2–5^. Generation of Cas9 knock-in mice have enabled *in vivo* cancer modelling and pooled genetic screening in primary immune cells, which has led to numerous discoveries of key tumor-growth drivers and immune regulators^6–9^. While Cas9 has been widely used since the inception of CRISPR editing era, the emergence of Cas12a presents distinct advantages, particularly in multiplexed gene editing^10, 11^. Apart from its DNase activity, Cas12a possesses RNase activity that allows generation of mature CRISPR RNAs (crRNAs) from an array of concatenated crRNAs by cleaving RNAs at the direct repeat (DR) consensus sequences^10, 11^. This unique capability of multiplex gene perturbation renders it an ideal candidate for elucidating intricate gene interactions such as epistasis, redundancy, synergy, and antagonism.

However, the large size of Cas12a (1,200 to 1,500 amino acids long) family proteins pose a substantial obstacle in terms of delivery and stable expression of CRISPR-Cas12a system by viral vectors, especially in primary cells^12^. Thus, we reasoned that developing Cas12a-knock-in mice would streamline the process of primary cell genome engineering and Cas12a-based CRISPR screening. We have developed LbCas12a-knock-in mice and the retroviral guide delivery system, which together demonstrated successful genome editing in primary murine immune cells^13^ (LbCas12a mice bioRxiv preprint, related manuscript). An engineered version of AsCas12a, enAsCas12a (with substitutions E174R/S542R/K548R) and its high-fidelity version enAsCas12a-HF1 (with substitutions E174R/N282A/S542R/K548R), were demonstrated to have expanded PAM sequence and enhanced multiplexed gene editing efficiency^14^. Therefore, developing mouse model with stable enAsCas12a-HF1 knock-in holds promising potential for efficient *in vivo* and *ex vivo* multiplexed genome engineering, particularly in primary murine cells.

Here, we generated both Cre-dependent, conditional LSL-enAsCas12a-HF1 and constitutive enAsCas12a-HF1 *Rosa26* locus knock-in mice, to facilitate efficient *in vivo* and *ex vivo* multiplexed genome engineering. These mice are compatible with multiple delivery vehicles including LNP-RNA, adeno-associated virus (AAV), and retroviral vectors. By delivering crRNAs using LNP into the constitutive enAsCas12a-HF1 mice, we achieved functional knockdown of the transthyretin (TTR) protein, a misfolded form of which leads to life-threatening transthyretin amyloidosis^15, 16^. We demonstrated efficient quadruplex gene knockout *in vivo* using a single AAV vector simultaneously targeting murine *Trp53, Apc, Pten*, and *Rb1* simultaneously, resulting in rapid induction of salivary gland squamous cell carcinoma and lung adenocarcinoma. Furthermore, we showed its capability of *ex vivo* multiplexed gene perturbation in primary immune cells. Finally, we demonstrated the modularity of LSL-enAsCas12a-HF1 mice by integrating with a CRISPR activation (CRISPRa) transgenic mouse line (dCas9-SPH) ^17^ to establish a simultaneous dual gene activation and knockout (DAKO) system. The enAsCas12a transgene was also independently knocked into a different safe harbor locus *H11* for creation of H11-enAsCas12a mice, which demonstrated utility in modeling of oncogene-negative lung adenocarcinoma, small-cell lung cancer ^18^ (H11-enAsCas12a mice bioRxiv preprint, related manuscript). The two species of Cas12a (LbCas12a and AsCas12a), as well as different knock-in strains, also offer multiple choices as expanded Cas12a-based gene editing technology applications.

## Results

### Generation of LSL-enAsCas12a-HF1 and enAsCas12a-HF1 mice

To generate Cre-dependent conditional LSL-enAsCas12a-HF1 knock-in mice, we cloned the enAsCas12a-HF1, tagged with MycTag and enhanced GFP (eGFP), into the Ai9 Rosa26-targeting construct. The construct contained Rosa26 homology arms for transgene knock-in between exon 1 and exon 2 of the *Rosa26* locus. The expression of enAsCas12a-HF1 was controlled by a CAG promoter and interrupted by a LoxP-3x PolyA Stop-LoxP (LSL) cassette upstream of the enAsCas12a-HF1 transgene, which allowed conditional enAsCas12a-HF1 expression by Cre recombinase (**Methods**) (**Fig. 1a**).

**Figure 1.**
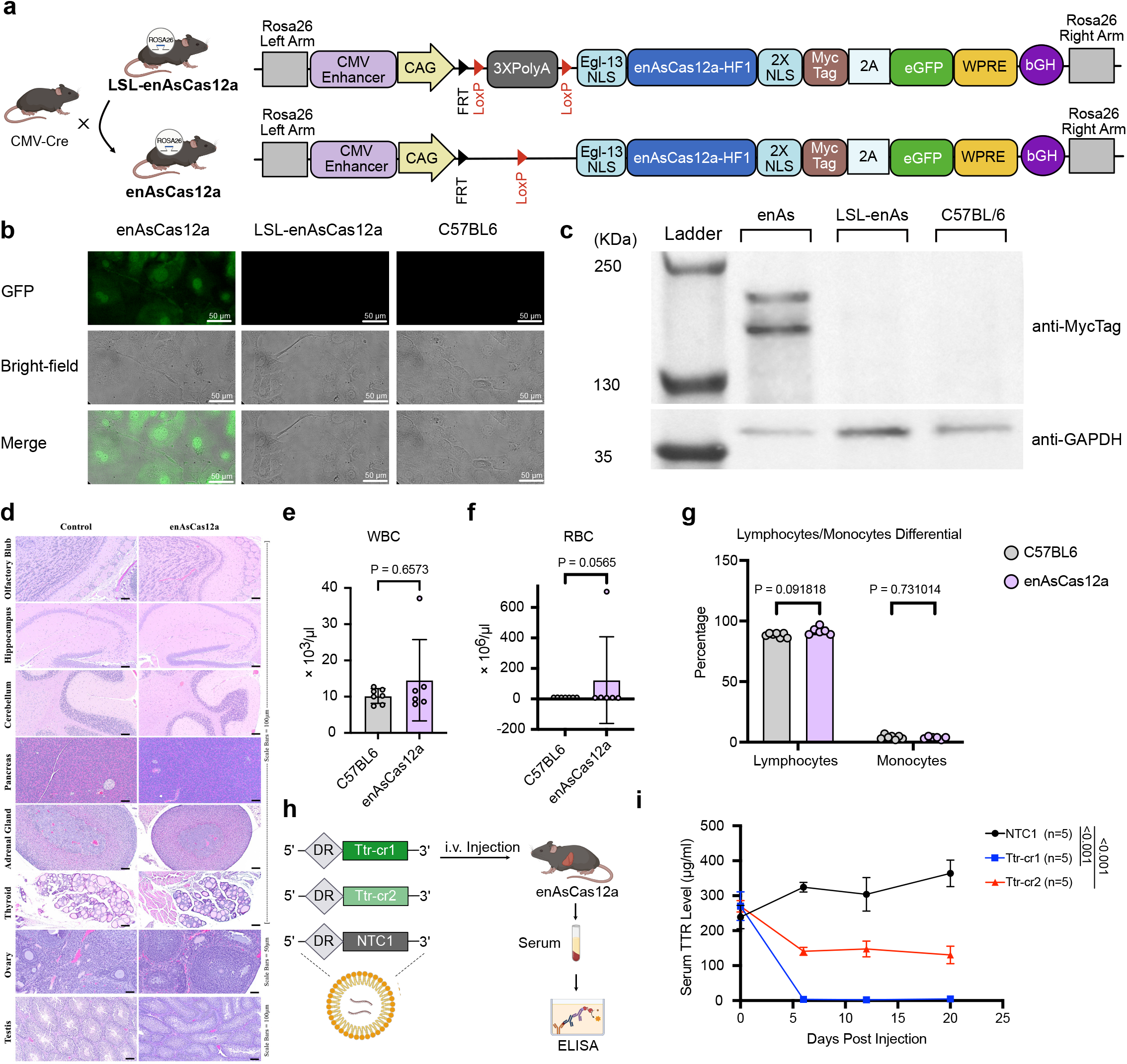
Generation of LSL-enAsCas12a-HF1 Mice and Liver Targeting with LNP-crRNA. a) Schematic showing the DNA sequence that was knocked into the Rosa26 locus. The backbone is Ai9 Rosa26 targeting vector. enAsCas12a-HF1, labeled with Myc tag and enhanced GFP, is expressed by CAG promoter. LoxP-(Stop)3XPolyA-LoxP (LSL) allows Cre-dependent conditional expression of enAsCas12a-HF1 protein. Egl-13 nuclear localization signal (NLS) was added at the N-terminus and nucleoplasmin NLS and c-Myc NLS signal were added at the C-terminus of enAsCas12a-HF. LSL-enAsCas12a-HF1 mice were then crossed with CMV-Cre mice to excise the LSL cassette to generate constitutive enAsCas12a-HF1 mice. b) Widefield fluorescent microscopy illustrating the expression of enAsCas12a-HF1-eGFP only in constitutive enAsCas12a-HF1 mouse, but not in conditional LSL-enAsCas12a-HF1 mouse or parental C57BL/6 mouse. c) Western blot showing the expression of enAsCas12a-HF1-MycTag protein in enAsCas12a-HF1 mouse. enAsCas12a-HF1-MycTag protein was not detected in protein lysate from LSL-enAsCas12a-HF1 and C57B/L6 mouse. Anti-MycTag antibody was used to detect enAsCas12a-HF1 protein and GAPDH was used as internal control. d) Representative histology of major organs from control mice (C57BL/6) and constitutive enAsCas12a-HF1 mice. e) White Blood Cell (WBC) count for constitutive enAsCas12a-HF1 mice and C57BL/6 mice. Mann-Whitney U Test was used to assess significance. For bar plot, data are shown as mean ± s.e.m. Exact *P* values are labeled. For both groups, n = 6. f) Red Blood Cell (RBC) count for constitutive enAsCas12a-HF1 mice and C57BL/6 mice. Mann-Whitney U Test was used to assess significance. For bar plot, data are shown as mean ± s.e.m. Exact *P* values are labeled. For both groups, n = 6 biological replicates. g) Comparison of lymphocytes/Monocytes differential between constitutive enAsCas12a-HF1 mice with C57BL/6 mice. Mann-Whitney U Test was used to assess significance. For bar plot, data are shown as mean ± s.e.m. Exact *P* values are labeled. For both groups, n = 6 biological replicates. h) Schematic showing the packaging and delivery of crRNA using lipid nanoparticle (LNP) to knockout liver protein TTR. Non-targeting control crRNA 1 (NTC1) were packaged as control in the same batch. Constitutive enAsCas12a-HF1 mice were intravenously (i.v.) injected with LNP-crRNA. Serum samples were harvested for ELISA to measure and monitor the TTR protein level in serum. i) Serum TTR level (µg/ml) from samples collected at Day0 (prior to injection), Day6, Day12, and Day20 via retro-orbital (RO) blood draw post injection. The serum TTR level was measured by ELISA. Two independent guides (TTRcr1 and TTRcr2) targeting murine *Ttr* gene were compared to non-targeting control crRNA (NTC1). Two-way ANOVA with Dunnett’s multiple comparisons test was used to assess significance. Data are shown as mean ± s.e.m. *P* values are labeled. For all groups, n = 5 biological replicates.

It was postulated that the suboptimal gene editing efficiency of Cas12a was in part due to the present of two nuclear export sequences in its conserved RuvC-II domain^19^. Multiple studies showed that increasing the number of SV40 nuclear localization signals (NLSs) on Cas12a augments its endonuclease activity in both zebra fish and mammalian cells^19, 20^. Cas12a with a combination of different NLSs was shown to outperform that with the same type of NLSs, while minimizing the off-target potential^21^. Furthermore, the position of these NLSs on the protein had drastic impact on nuclear localization^22^. Taking these factors into consideration, we placed the Egl-13 NLS on the N-terminus, and the nucleoplasmin NLS and c-Myc NLS signal on the C-terminus of enAsCas12a-HF1 to maintain higher concentration of enAsCas12a-HF1 in the nucleus (**Methods**) (**Fig. 1a**).

We then co-injected SpCas9_Rosa26-sgRNA ribonucleoprotein (RNP) with a linearized enAsCas12a-HF1-Rosa26-targeting construct to generate the Rosa26-knock-in mouse strain LSL-enAsCas12a-HF1, in the C57BL/6 background for its broad utility in immunology and immune-oncology fields. The founding generation (F0) animals’ genotypes were verified by polymerase chain reaction (PCR) and Sanger sequencing. The F0 animals were backcrossed to C57BL/6 background to establish the F1 generation. The F1 heterozygotes were crossed to each other to achieve homozygosity (F2 and beyond). (**Methods**) (**Fig. 1a**).

LSL-enAsCas12a-HF1 mice were then crossed with the CMV-Cre mice^23^ to generate constitutive enAsCas12a-HF1 mice (**Methods**) (**Fig. 1a**). To confirm successful generation of constitutive enAsCas12a-HF1 mice, we isolated primary ear fibroblasts from enAsCas12a-HF1, LSL-enAsCas12a-HF1, and C57BL/6 mice, and determined the protein expression of enAsCas12a-HF1 by widefield fluorescence microscopy and Western blot. We observed that GFP signal was detected only in enAsCas12a-HF1 fibroblasts but not in LSL-enAsCas12a-HF1 or C57BL/6 fibroblasts, indicating enAsCas12a-HF1-2A-eGFP was only detectable in the constitutive enAsCas12a-HF1 mice (**Fig. 1b**), implying a tight expression control by the LSL allele. In concordance with the microscopy result, Western blot analysis using an anti-MycTag antibody on cell lysate of the ear fibroblasts showed enAsCas12a-HF1-NLS-Myc protein expression only in constitutive enAsCas12a-HF1 mice (**Fig. 1c**). These results demonstrated the successful generation of the two knock-in mouse strains and confirmed the tightly controlled expression of Cas12a protein in LSL-enAsCas12a-HF1 mice at least by measurable reporter (eGFP) and tag (Myc).

In contrast to Cas9, Cas12a family proteins were reported to have a unique “*trans*-cleavage” activity, by which Cas12a indiscriminately degrades single-stranded DNA (ssDNA) upon activation by RNA-guided DNA binding^24^. Thus, it is critical to characterize the impact of constitutive and prolonged enAsCas12a-HF1 expression on the physiological traits or pathological effects of the knock-in mice. The constitutive enAsCas12a-HF1 mice did not show any noticeable difference from wild-type (WT) mice in terms of fertility, morphology, and were able to breed to and maintain homozygosity. We also sent a batch of enAsCas12a-HF1 mice for professional and blinded necropsy and pathology analysis, together with the background strain, C57BL/6 mice, at similar age, housed in the same condition. Hematoxylin and eosin (H&E) staining showed no significant findings in the morphological differences between the WT mice and the constitutive enAsCas12a-HF1 mice, nor any detectable pathological signs in either strain, for all tissues and organs examined, including the brain, pancreas, adrenal gland, thyroid, and reproductive organs from both sexes (ovary/testis) (**Fig. 1d**). Complete blood count (CBC) analysis demonstrated negligible differences in representative values like white blood cell count, red blood cell count, and lymphocytes/monocytes differential (**Fig. 1e-g**). These results together suggested that there is no noticeable toxicity caused by constitutive enAsCas12a-HF1 expression.

### *In vivo* liver gene targeting mediated by LNP-crRNA

Viral vectors are widely used in the delivery of CRISPR systems for *in vivo* genome editing and for gene therapy^6, 25, 26^. However, these viral vectors might elicit adaptive immune response which limits their therapeutic efficacy^27^. Using them as an *in vivo* delivery method has also raised safety concerns since the death of a Duchenne muscular dystrophy patient after high-dose AAV gene therapy^28^. Meanwhile, LNP, a non-viral delivery method, has been proven safe and effective preclinically and clinically, both for mRNA delivery as COVID-19 vaccines ^29, 30^, and for CRISPR gene therapy in the liver^31^. To demonstrate this application in constitutive enAsCas12a-HF1 mice and to test this in a disease related model, we delivered LNP-crRNA through intravenous (i.v.) injection (**Fig. 1h**) to target the murine *Ttr* gene, whose human homolog is a known therapeutic target of transthyretin amyloidosis^15, 16^. We parallelly encapsulated two independent guides (*Ttr*-cr1, and *Ttr*-cr2) targeting *Ttr* gene, together with a non-targeting control NTC1 (**Fig. 1h**). Both*Ttr*-cr1, and *Ttr*-cr2 induced rapid and significant serum TTR protein knockdown starting from day 6 post injection (**Fig. 1i**). Notably, the decrease in serum TTR levels remained stable during the 20 days’ monitoring period (**Fig. 1i**). We observed near 100% knockdown of serum TTR level by *Ttr*-cr1 relative to NTC1 (**Fig. 1i**). *Ttr*-cr2, which ranked lower in the CRISPick algorithm^32^, also induced near 50% knockdown of serum TTR (**Fig. 1i**). These results demonstrated the highly efficient *in vivo*, therapeutically relevant gene targeting capability of the enAsCas12a-HF1 mice when coupled with the LNP-crRNA delivery system, which provides broad applicability for direct *in vivo* targeting and delivery studies in disease models.

### LSL-enAsCas12a-HF1 mice enabled multiplexed *in vivo* gene editing for autochthonous cancer modeling

Genetically engineered mouse models (GEMMs) allow autochthonous cancer modeling – inducing tumors from normal cells *de novo* within the intact organism. Compared with xenograft or syngeneic models that rely on cell line transplantation, autochthonous models bear a more *bona fide* resemblance of the human oncogenesis process and tumor microenvironment, which are critical for the study of cancer biology, immune surveillance and therapeutic response^33^. The conventional method of creating GEMMs, which has conditional knockout alleles for each of the tumor suppressors, is a tedious process^34^. The use of CRISPR-mediated genome engineering, especially the generation of Cas9 knock-in mice and associated viral delivery methods, substantially simplified autochthonous cancer modeling process, and could be scaled up to systematically screen for drivers of tumor oncogenesis and progression in different cancer types^6–8, 35–37^. However, two bottlenecks severely restrict the effectiveness of Cas9 systems for tumor modeling– 1) lack of multiplex gene editing capability for Cas9, and 2) packaging size constraint of AAV vector. These limitations effectively hinder Cas9 systems from perturbing scaling numbers of genes encoding cancer drivers or regulators. On the other hand, Cas12a has been shown to have high multiplexing capability in gene editing using a crRNA array encoding multiple crRNAs targeting multiple genes in single vector^38^. We reasoned that LSL-enAsCas12a-HF1 mice would serve as an efficient tool for more adaptable and precise autochthonous cancer modeling.

To experimentally test this, we selected from the pan-cancer database MSK-IMPACT 6 frequently mutated tumor suppressor genes (TSGs): *TP53*, *APC*, *PTEN*, *RB1*, *SMAD4*, and *STK11* in a pan-cancer analysis^39, 40^ (**Fig. 2a**). After pairwise co-occurrence (CO) and mutual exclusivity (ME) analysis, we picked *TP53*, *APC*, *PTEN*, and *RB1,* all of which co-occurred pair-wisely across multiple types of human cancers (collectively in a pan-cancer manner)^39, 40^ (**Fig. 2b**). To target murine *Trp53, Pten, Apc*, and *Rb1* simultaneously using a single AAV vector, we cloned a concatenated RNA array (crTSG) containing four crRNAs (crTrp53, crPten, crApc, and crRb1), separated by enAsCas12a direct repeats (DRs), driven by a single U6 promoter, together with a Cre recombinase expressed under a constitutive promoter EFS to activate enAsCas12a-HF1-eGFP expression (**Fig. 2c**). When intravenously injected with this AAV-crTSG-Cre, 100% (6/6) mice developed aggressive palpable head and neck cancer within a month of the single dose, whereas none (0/6) of the AAV-vector (that also expresses Cre) injected mice showed any noticeable carcinogenesis (**Fig. 2d**). Because the 2A-eGFP is co-cistronically encoded with the enAsCas12a-HF1 transgene, AAV-infected cells in the LSL-enAsCas12a-HF1-2A-eGFP mice would emit green fluorescence as Cre recombinase activates enAsCas12a-HF1-eGFP expression. Significantly stronger GFP signals were indeed detected in head and neck tumor samples compared to those in other organs, due to the rapid growth of cancer cells (**Fig. 2d, f**). Of note, other than the tumor, as expected the liver has relatively higher signal of eGFP reporter as compared to other organs examined (spleen and brain) (**Fig. 2d, 2e, 2f**).

**Figure 2.**
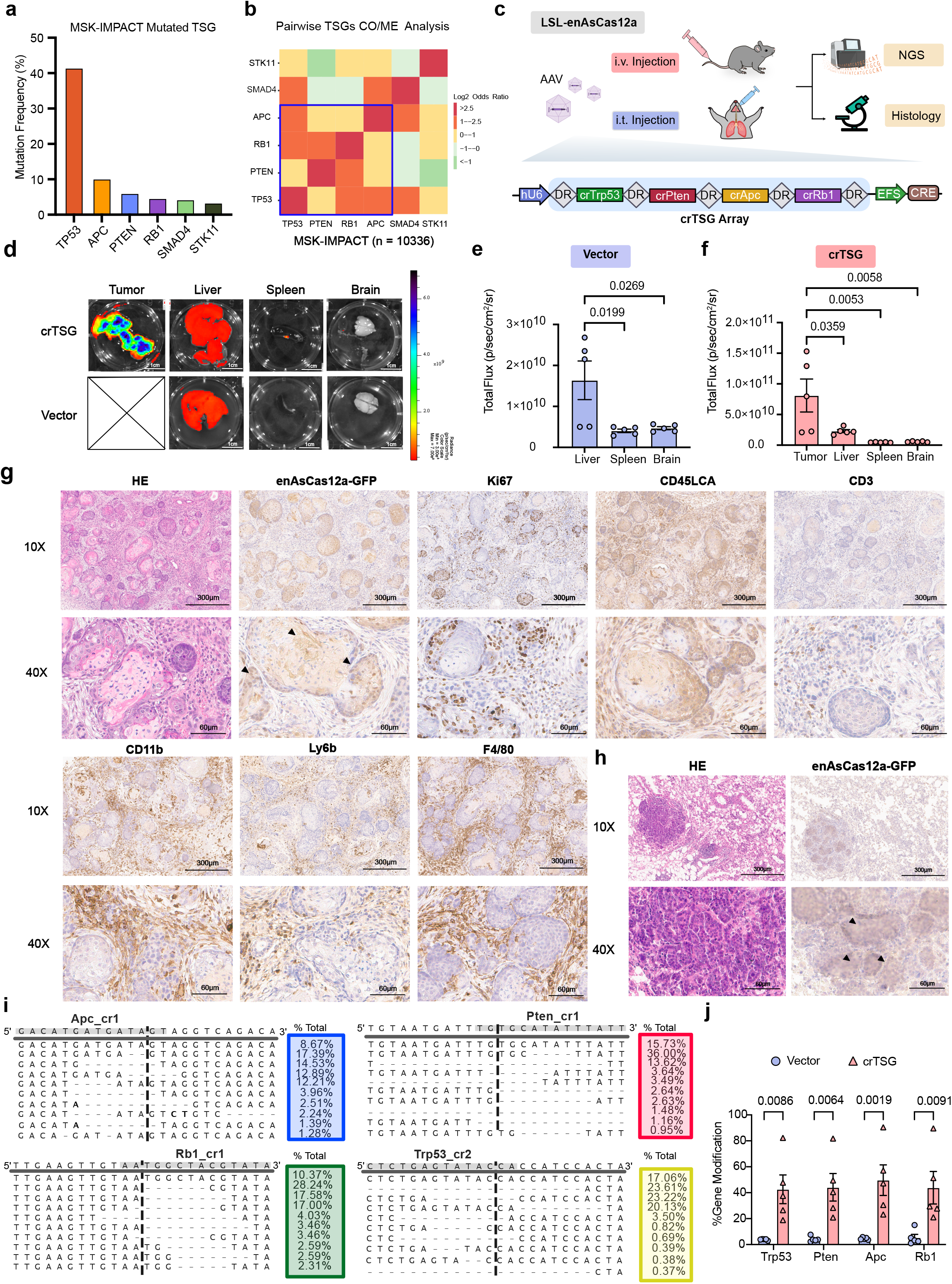
Tumorigenesis Induced by AAV-Mediated *in vivo* Gene Editing. a) Mutation frequency of some common tumor suppressor genes (TSGs), *TP53, APC, PTEN, RB1, SMAD4*, and *STK11* from a pan-cancer database (MSK-IMPACT project)^39, 40^. b) Pairwise co-occurrence (CO) and mutually exclusivity (ME) analysis for TSGs, *TP53, APC, PTEN, RB1, SMAD4*, and *STK11*. Positive Log2(Odds Ratio) indicated co-occurrence while negative Log2(Odds Ratio) indicated mutual exclusivity^39, 40^. We identified *TP53, APC, PTEN,* and *RB1* as a group of co-occur genes out of the 6 TSGs (surrounded by blue lines). *SMAD4* was excluded due to mutual exclusivity with *PTEN, RB1* and *STK11*. *STK11* was excluded because it was mutually exclusive with *PTEN* and *SMAD4*. c) Schematic showing part of the AAV construct used for tumor induction, which contained a crRNA expression array (crTSG) that had 4 guides targeting *Trp53, Pten, Apc, and Rb1*, and a Cre expression cassette for inducing enAsCas12a-HF1 expression. After production and purification, AAV-crTSG and AAV-vector (as negative control) were either intravenously (i.v.) or intratracheally (i.t.) injected into LSL- enAsCas12a-HF1 mice. Tumor and major organs were isolated for Next Generation Sequencing (NGS) and histology analysis. d) Representative IVIS Spectrum images detecting GFP signal indicated by total radiance (p s^−1^ cm^−2^ sr^−1^) for tumor and major organs. Top row being representative AAV-crTSG i.v. injected mouse and the bottom row being representative AAV-vector i.v. injected mouse. e) Quantification of GFP signals indicated by total radiance (p s^−1^ cm^−2^ sr^−1^) for AAV-vector group. One-way ANOVA with Tukey’s multiple comparisons test was used to assess significance. For bar plot, data are shown as mean ± s.e.m. Exact *P* values are labeled. For all organs, n = 5 biological replicates. f) Quantification of GFP signals indicated by total radiance (p s^−1^ cm^−2^ sr^−1^) for AAV-crTSG group. One-way ANOVA with Tukey’s multiple comparisons test was used to assess significance. For bar plot, data are shown as mean ± s.e.m. Exact *P* values are labeled. For all organs, n = 5 biological replicates. g) Representative hematoxylin & eosin (H&E) staining and immunohistochemistry (IHC) staining on AAV-induced salivary gland squamous cell carcinoma (SCC) FFPE samples. For IHC, tumor samples were stained with GFP (enAsCas12a-HF1-eGFP), Ki67 (proliferation marker), CD45LCA (immune cells marker), CD3(T cell marker), CD11b (myeloid lineage marker), Ly6b (neutrophile marker), and F4/80 (macrophage marker). For 10X images, scale bar = 300µm. For 40X images, scale bar = 60µm. h) Representative hematoxylin & eosin (H&E) staining and immunohistochemistry (IHC) staining on AAV-induced lung adenocarcinoma FFPE samples. For IHC, tumor samples were stained with GFP (enAsCas12a-HF1-eGFP). For 10X images, scale bar = 300µm. For 40X images, scale bar = 60µm. i) Representative allele frequency plots of the crRNAs targeted sites of *Trp53, Apc, Pten*, and *Rb1* demonstrating the gene modification generated in squamous cell carcinoma (SCC) samples. j) Percent gene modification quantified for the 4 targeted genes, *Trp53, Apc, Pten*, and *Rb1,* by NGS for SCC samples. Two-way ANOVA with Šídák’s multiple comparisons test was used to assess significance. Data are shown as mean ± s.e.m. *P* values are labeled. For all genes, n = 5 biological replicates.

To examine the AAV-crTSG-Cre induced head and neck cancer model in these mice, we performed endpoint histology analysis, which revealed characteristic squamous cell carcinoma (SCC) features (abundant eosinophilic cytoplasm and variable keratinization)^41^. Diagnosed analysis of the SCC and its adjacent tissues by a professional pathologist (C. L.) suggested that it originated from salivary glands (**Fig. 2g**). Consistent with the GFP signals detected by IVIS spectrum, the tumor sections showed enAsCas12a-HF1-GFP expression in immunohistochemistry (IHC) analysis, suggesting the tumor induction by the AAV-crTSG-Cre mediated gene editing (**Fig. 2g**). We observed that most of the SCC tumor cells were Ki67 positive, indicating a highly proliferative and aggressive nature of the cancer cells in this model (**Fig. 2g**). We also observed immune cells infiltration (marker CD45LCA), with higher infiltration by myeloid lineage immune cells (marker CD11b), such as neutrophiles (marker Ly6b) and macrophages (marker F4/80), than by T cells (marker CD3) (**Fig. 2g**). Intratracheal injection of the same AAV into the lung induced multifocal adenocarcinoma in lung, which also showed enAsCas12a-HF1-GFP expression as detected by IHC (**Fig. 2h**). Next Generation Sequencing (NGS) of the SCC samples showed significant levels of gene modification for the 4 targeted TSGs (*Trp53*, *Apc*, *Pten*, and *Rb1*) (on average approximately 50%, varying between genes); most of the modifications were deletions, centered around the predicted enAsCas12a crRNA cutting sites for these genes (**Fig. 2i, j**). Together, these results showcased the robustness of using LSL-enAsCas12a-HF1 mice for *in vivo* multiplexed genome engineering, which empowered autochthonous cancer modeling.

### LSL-enAsCas12a-HF1 mice enabled multiplexed *ex vivo* genome engineering in mouse primary immune cells

To evaluate the capability of the LSL-enAsCas12a-HF1 mice for multiplexed gene engineering in primary immune cells, we tested double-knockout gene editing in bone marrow derived dendritic cells (BMDCs) and CD8 T cells (**Fig. 3a**). We first generated a retroviral system (pRetro-hU6-AsDR-BbsI/crRNA-EFS-Cre-mScarlet) for *ex vivo* crRNA delivery to facilitate efficient gene editing with LSL-enAsCas12a-HF1 mice. The vector contained a single U6 promoter to express the enAsCas12a-HF1 crRNA array, which can be directly cloned into a dual BbsI digestion site downstream of the AsCas12a direct repeat (AsDR) (**Fig. 3a**). We also incorporated a Cre expression cassette for inducing enAsCas12a-HF1 expression in primary immune cells from the LSL-enAsCas12a-HF1 mice, and a mScarlet reporter for monitoring transduction efficiency (**Fig. 3a**). CD8 T cells and BMDCs were harvested from spleen and bone marrow, followed by spin infection and subsequent FACS analysis and sorting (**Fig. 3a**). CD11c/CD11b and CD8a were used as markers for BMDCs and CD8 T cells, respectively.

**Figure 3.**
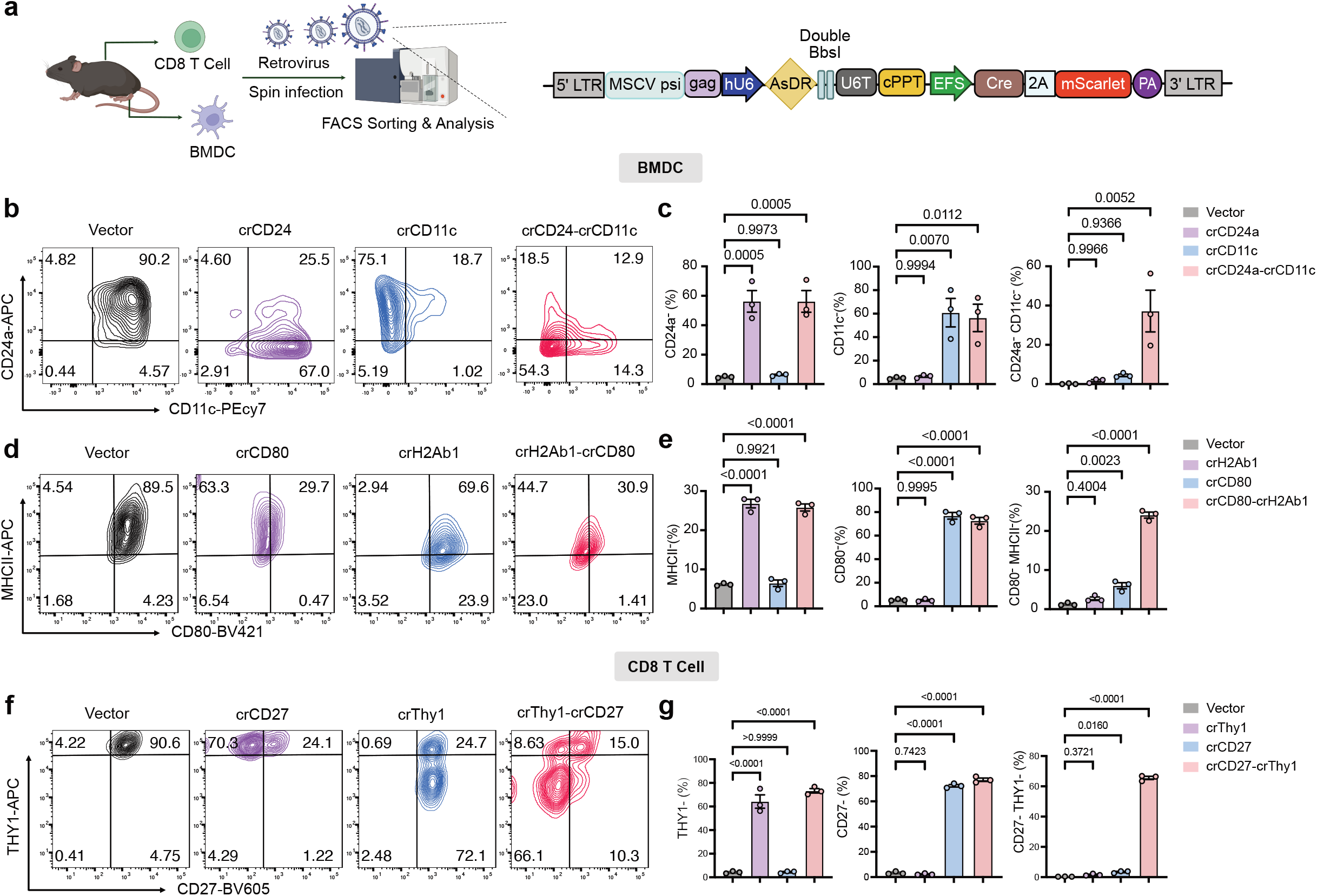
Multiplexed immune cell gene editing with LSL-enAsCas12a-HF1 mice. a) Schematic showing the retroviral vector design and *ex vivo* workflow for multiplexed gene editing in primary immune cells. CrRNA array was expressed by human U6 promoter (hU6). Cre and mScarlet expression was driven by EFS promoter to induce enAsCas12a-HF1 expression and to facilitate sorting for infected cells respectively. CD8 T cells and BMDCs were isolated from the spleen and bone marrow of LSL-enAsCas12a-HF1 mice, followed by *ex vivo* culture and retroviral infection. FACS was used to analyze the efficiency of single-cell level double knockout and enrich for infected cells by sorting. b) Representative contour plots of mScarlet^+^ BMDCs showing the *CD24* expression in relation of the *CD11c* expression. Anti-CD24a-APC and anti-CD11c-PE/Cy7 were used in the flow staining. Double knockout crRNA containing both crCD24 and crCD11c was compared with single crRNA (crCD24 and crCD11c) and vector control. c) Quantification of CD24a^-^, CD11c^-^, and CD24a^-^;CD11c^-^ percentage with respect to mScarlet^+^ BMDCs. One-way ANOVA with Tukey’s multiple comparisons test was used to assess significance. For bar plot, data are shown as mean ± s.e.m. Exact *P* values are labeled. For all groups, n = 3 biological replicates. d) Representative contour plots of mScarlet^+^ BMDCs showing the *MHCII* expression in relation of the *CD80* expression. Anti-MHCII-APC and anti-CD80-BV421 were used in the flow staining. Double knockout crRNA containing both crH2Ab1 and crCD80 was compared with single crRNA (crH2Ab1 and crCD80) and vector control. e) Quantification of MHCII^-^, CD80^-^, and MHCII^-^;CD80^-^ percentage with respect to mScarlet^+^ BMDCs. One-way ANOVA with Tukey’s multiple comparisons test was used to assess significance. For bar plot, data are shown as mean ± s.e.m. Exact *P* values are labeled. For all groups, n = 3 biological replicates. f) Representative contour plots of mScarlet^+^ CD8 T cells showing the *Thy1* expression in relation of the *CD27* expression. Anti-THY1-APC and anti-CD27-BV605 were used in the flow staining. Double knockout crRNA containing both crThy1 and crCD27 was compared with single crRNA (crThy1 and crCD27) and vector control. g) Quantification of THY1^-^, CD27^-^, and THY1^-^;CD27^-^ percentage with respect to mScarlet^+^ CD8 T cells. One-way ANOVA with Tukey’s multiple comparisons test was used to assess significance. For bar plot, data are shown as mean ± s.e.m. Exact *P* values are labeled. For all groups, n = 3 biological replicates.

To facilitate the detection of protein-level knockdown at the single cell resolution by flow cytometry, we focused on highly expressed surface markers. In BMDCs, we first tested two sets of dual gene targeting / double knockouts (DKOs), CD24-CD11c and CD80-H2Ab1, each of which was compared to their respective single knockouts and the vector control. Flow cytometry data showed that crCD24-crCD11c induced an average of 56.2% cell-population-level protein knockdown of CD24, and an average of 56.4% cell-population-level knockdown of CD11c, which was comparable to the efficiency of the single knockout controls (**Fig. 3b, 3c**). Notably, crCD24-crCD11c generated efficient CD24 and CD11c double knockdown at the single cell level, with an average of 37.2% CD24^-^;CD11c^-^ double negative population; whereas single knockout controls, crCD24 and crCD11c, did not result in a noticeable double negative population (**Fig. 3b**, **Fig. 3c**). This suggests that a fraction of the transduced cells have achieved double knockdown at the single-cell level generated by the crCD24-crCD11c crRNA array in BMDCs. Similarly, crCD80-crH2Ab1 induced 72.2% knockdown of CD80 and 25.8% knockdown of MHCII, which was comparable to the efficiency of the single knockout controls (**Fig. 3d**, **Fig. 3e**). Importantly, an average of 24% of the cells showed CD80^-^;MHCII^-^ double negative in the crCD80-crH2Ab1 group, again suggesting single-cell-level double knockdown for a fraction of the transduced cells. (**Fig. 3d**, **Fig. 3e**).

Efficient gene editing was also achieved in CD8 T cells. Single knockout controls, crThy1 and crCD27, resulted in averages of 64.2% and 72.3% cell-population-level protein knockdown of THY1 and CD27, respectively (**Fig. 3f, 3g**). The DKO construct, crThy1-crCD27, induced similar levels of cell-population-level protein knockdown of both proteins as compared to the single knockout controls (**Fig. 3f, 3g**). Importantly, only the DKO construct resulted in an average of 65.6% THY1^-^;CD27^-^ population (**Fig. 3f**, **Fig. 3g**), suggesting that 65.6% of the transduced cells achieved double knockdown at the single-cell level by the crThy1-crCD27 DKO crRNA array in CD8 T cells. Collectively, these results demonstrated the multiplexed immune cell genome engineering capability of the Cas12a knock-in mice.

### Development of a simultaneous dual gene activation and knockout (DAKO) system using the LSL-enAsCas12a-HF1 mice

Attempts have been made to systematically studying both the positive and negative regulators for the same biological processes^42, 43^. These studies have relied on either conduction of two independent genetic manipulations (one loss-of-function and one gain-of-function), or relying on directions (upregulation and downregulation) of a single type of perturbation via data analysis ^42, 43^. However, these methods are indirect. To date, there remains a lack of an efficient tool for simultaneous gene activation and knockout, in particular within a single cell. To take on this challenge, we developed a DAKO system by crossing the LSL-enAsCas12a-HF1 mice with the dCas9-SPH mice (dCas9 fused with activation domain SunTag-p65-HSF1 for potent transcriptional activation)^17^ and selecting for double positive progenies by genotyping to generate the LSL-enAsCas12a-HF1;dCas9-SPH DAKO mice (**Fig. 4a**, **Fig. 4b**). We also designed a Cas9-Cas12a fusion guide-RNA system that expresses both types of CRISPR guide RNAs (Cas9 sgRNA and Cas12a crRNA) in a string: Cas9 sgRNA was concatenated with Cas12a crRNA, separated by AsCas12a DR sequence, for simultaneously expression in the same cells (**Fig. 4b**). We hypothesized that this guide RNA chimera would be cleaved by enAsCas12a-HF1 protein, upstream of AsDR, into a mature Cas9-sgRNA and a mature Cas12a crRNA, which would then direct the gene activation by dCas9-SPH and gene knockout by enAsCas12a-HF1 (**Fig. 4b**).

**Figure 4.**
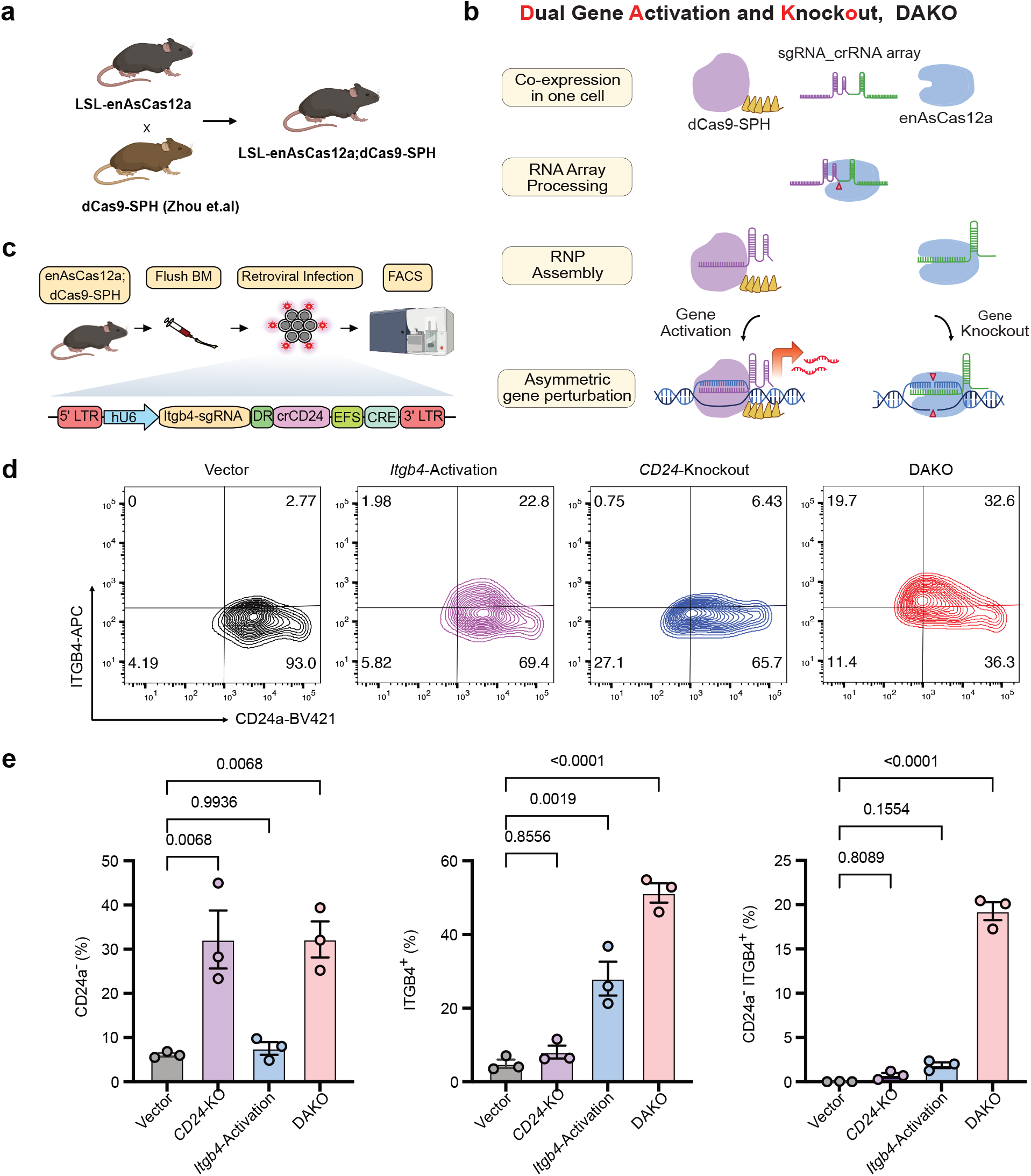
Dual gene activation and knockout (DAKO) with LSL-enAsCas12a-HF1;dCas9-SPH double transgenic mice. a) Schematic showing the breeding strategy of LSL-enAsCas12a-HF1;dCas9-SPH mice. b) Schematic demonstrating the mechanism of the simultaneous Dual Activation and Knockout (DAKO) system. A dCas9-sgRNA was concatenated with Cas12a Direct Repeat (DR) and crRNA, which was then processed and cleaved into mature dCas9-sgRNA and Cas12a-crRNA by Cas12a protein, upon delivery into the same cell. The mature dCas9-sgRNA and Cas12a-crRNA then assembled with corresponding protein to mediate gene activation and gene knockout respectively. RNP: ribonucleoprotein. c) Schematic illustrating the retroviral construct and the workflow of *ex vivo* of DAKO validation in primary BMDCs. Human U6 promoter (hU6) drove the expression of a Cas9 activating guide for *Itgb4* (Itgb4-sgRNA) was concatenated with Cas12a direct repeat (DR) and crRNA targeting *CD24* (crCD24). A Cre expression cassette was used to induce enAsCas12a-HF1 expression. BM: bone marrow. d) Representative contour plots of mScarlet^+^ BMDCs showing the I*tgb4* expression in relation of the *CD24a* expression. Anti-ITGB4-APC and anti-CD24a-BV421 were used in the flow staining. DAKO-crRNA containing both crItgb4 and crCD24 was compared with single crRNA (Itgb4-sgRNA and crCD24) and vector control. e) Quantification of CD24a^-^, ITGB4^+^, and CD24a^-^;Itgb4^+^ percentage with respect to mScarlet^+^ BMDCs. One-way ANOVA with Tukey’s multiple comparisons test was used to assess significance. For bar plot, data are shown as mean ± s.e.m. Exact *P* values are labeled. For all groups, n = 3 biological replicates.

To validate this system in primary immune cells, we isolated BMDCs from LSL-enAsCas12a-HF1;dCas9-SPH DAKO mice and infected them with retrovirus expressing a DAKO fusion guide cassette containing an Itgb4-sgRNA concatenated with an AsDR-crCD24, alongside a constitutively active EFS-Cre cassette (sgItgb4-crCD24-Cre vector) (**Fig. 4c**). We additionally included the vector, single activation (Itgb4-sgRNA), and single knockout (crCD24) groups in parallel as controls. Flow cytometry analysis showed that the sgItgb4-crCD24-Cre DAKO targeting resulted in a comparable cell population level protein knockdown of CD24 knockdown as that of the single knockout control (crCD24-Cre) (average 32.2% CD24-negative population in both groups) (**Fig. 4d**, **Fig. 4e**). The sgItgb4-crCD24-Cre DAKO also resulted in a cell-population-level protein upregulation of ITGB4 (average 51% ITGB4-positive population in DAKO) as compared to that of the single gene CRISPRa targeting control (average 27% ITGB4-positive population in sgItgb4-Cre control) (**Fig. 4d**, **Fig. 4e**). Importantly, only in the DAKO group did we observe a significant level of anticipated dual-targeting population of cells with the correct direction of gene regulation (average of 19.3% CD24^-^;Itgb4^+^ cells in the sgItgb4-crCD24-Cre DAKO group, p < 0.0001), as compared to background levels in both the sgItgb4-Cre and the crCD24-Cre control groups (p > 0.1, not significant) (**Fig. 4d, 4e**). These results suggested that the DAKO system (enAsCas12a-HF1;dCas9-SPH DAKO mice plus the Cas9-sgRNA-Cas12a-crRNA chimeric fusion guides) could achieve efficient and simultaneous gene activation and knockout at the single cell level, enabling orthogonal dual gene targeting in immune cells.

## Discussion

Deconvolution of the complex gene regulation networks is fundamental for understanding basic biology to the pleiotropic nature of human disease. Thus, a versatile platform for multiplexed *in vivo* and *ex vivo* genome engineering is needed to systematically study the gene interactions. Here, we generated both conditional LSL-enAsCas12a-HF1 and constitutive enAsCas12a-HF1 mice to allow highly efficient genome engineering in primary cells. For compatible utilization of these strains, multiple delivery methods (LNP, AAV, and retrovirus) were also developed and applied in conjunction with these mice for efficient liver gene targeting, tumor modeling, and primary immune cell editing respectively. The liver *Ttr* gene was efficiently knocked out within a week using LNP-crRNA. Salivary gland squamous cell carcinoma was induced rapidly within a month by targeting 4 TSGs for simultaneous knockout. Efficient double knockout was demonstrated in both primary BMDCs and CD8 T cells. The DAKO system also showcased the asymmetric genetic manipulation in primary immune cells.

Our constitutive and conditional knock-in enAsCas12a-HF1 strains enable efficient *in vivo* and *ex vivo* multiplexed genome engineering and provide opportunities for utilization in diverse fields. These mice demonstrate editing efficiency comparable to that of Cas9 knock-in mice and have unique advantages of crRNA array based multiplex gene editing compared to the Cas9 mice. Additionally, these mice can facilitate rapid and seamless workflows for *in vivo* therapeutic gene targeting, disease / tumor modeling and primary immune cell engineering, as well as future multiplexed genetic manipulations at higher throughput by others in the field. The generation of Cas12a mice with other variations, such as a different species of the Cas12a^13^ (LbCas12a mice bioRxiv preprint, related manuscript), or knock-in into a different safe harbor locus^18^ (H11-enAsCas12a mice bioRxiv preprint, related manuscript), represents represent convergent collective development of broadly enabling Cas12a gene editing toolkits for applications in diverse fields.

## Methods

### Institutional Approval

The study has obtained regulatory approval from the relevant institutions. All activities involving recombinant DNA and biosafety were carried out in accordance with the guidelines set by the Yale Environment, Health, and Safety (EHS) Committee, following an approved protocol (Chen-rDNA 15-45; 18-45; 21-45). Furthermore, all animal-related work adhered to the guidelines established by the Yale University Institutional Animal Care and Use Committee (IACUC), following approved protocols (Chen 2018-20068; 2021-20068).

### Mouse Line Generation

LoxP-Stop-LoxP (LSL)-enAsCas12a-HF1 mice were produced through the pronuclear injection of a transgenic expression cassette consisting of IDT Alt-R HiFi Cas9 Nuclease V3 and a Rosa26-targeting crRNA (5’-ACTCCAGTCTTTCTAGAAGA-3’) (Chu et al., 2015). The Ai9 Rosa26-targeting vector was used to subclone a codon-optimized enAsCas12a-HF1 (with substitutions E174R/N282A/S542R/K548R) cDNA, resulting in the creation of the LSL-Egl_13-enAsCas12a_HF1-NLS-Myc-2A-eGFP transgenic expression cassette (Madisen et al., 2010; Sanjana et al., 2014). Unique sequencing and genotyping primers were designed using the NCBI Primer Blast tool, targeting the Mus musculus genome. Internal enAsCas12a-HF1 primers (forward: 5’-TTTCCACGTGCCTATCACACT-3’ and reverse: 5’-GCCCTTCAGGGCGATGTG-3’) were employed to confirm the presence of the enAsCas12a-HF1 transgene in knock-in mice. Rosa26 primers external to the Ai Rosa26-targeting vector were used for amplification and Sanger sequencing to verify the accurate integration of the transgenic expression cassette. Constitutive enAsCas12a-HF1 mice were generated by crossing LSL-enAsCas12a-HF1 mice with CMV-Cre mice (The Jackson Laboratory, Bar Harbor, ME). Additional mouse strains, including C57BL6/J (The Jackson Laboratory, Bar Harbor, ME), were employed for breeding and experimental purposes. All animals were housed in standard individually ventilated cages, with a light cycle of either 12 hours light and 12 hours dark or 13 hours light and 11 hours dark. The room temperature ranged from within 21-23°C, and the relative humidity was maintained between 40% and 60%.

### Mouse Primary Ear Fibroblast Cultures

Mouse ear tissue was harvested and sterilized by incubating in 70% EtOH for 3 mins. Primary fibroblast cultures were obtained by subjecting small pieces of mouse ear tissue to a 30-minute digestion with Collagenase/Dispase (Roche) at 37°C under agitation. The resulting supernatant was collected and washed with 2% FBS. After filtration through a 40μm filter, the cell suspensions were resuspended in DMEM media supplement with 10% FBS and 1% Pen/Strep.

### Widefield Fluorescence Microscopy

Primary ear fibroblast was first plated in 8 well glass bottom µ-slide (Ibidi), which was pre-coated with 2µg/cm^2^ poly-L-lysine (Sigma-Aldrich) according to manufacturer’s recommendations. Widefield fluorescent images of the mouse primary fibroblast cultures were taken using Leica DMi8 Widefield Fluorescence Microscope. The images were captured when the cells reached a confluency level of about 80%.

### Western Blot Analysis

Mouse primary ear fibroblast cultures mentioned above were washed with DPBS and collected using cell scrapper and lysis buffer, RIPA buffer (Boston BioProducts) supplemented with Halt ™ proteinase inhibitor cocktail (Thermo Fisher). Cells were lysed for 1hr in 4°C with agitation. Then, cell lysates were spun down at 4°C, 2000xg for 20 mins and protein-containing supernatants were collected. 4x Laemmli Sample Buffer (Bio-Rad, #1610747) with beta-mercaptoethanol (Sigma-Aldrich) was added to supernatant, followed by heat incubation at 95°C for 5 mins for denaturation. Protein concentrations were quantified using Pierce™ BCA Protein Assay kit (ThermoFisher, 23225) and normalized before proteins were loaded on 4∼20% Tris-Glycine gels for electrophoresis. Proteins on gel were transferred to 0.2μm nitrocellulose membranes. Membranes were blocked with 2.5% bovine serum albumin and stained with mouse anti-MycTag (Cell Signaling Technology, mAb#2278,1:1000), and 1:2000 rabbit anti-GAPDH (Thermo Fisher, MA1-16757, 1:2000) overnight at 4°C. The next day, membranes were incubated with secondary antibodies, anti-mouse IgG (Cell Signaling, 7076,1:5000) and anti-rabbit IgG (Cell Signaling, 7074,1:5000) for 1 h at room temperature. ECL prime western blotting detection reagents (Bio-Rad) were used for chemiluminescence detection and imaged using BioRad gel doc.

### Pathology analysis of enAsCas12a-HF1 mice

Nineteen mice: C57BL/J mice (4 male, 3 female) and 6 enAsCas12a-HF1 mice (4 male, 2 female), were submitted to the Comparative Pathology Research Pathology (CPR) Core, Yale University, Department of Comparative Medicine for necropsy, histology, and comprehensive histopathologic analysis blind to experimental manipulation. The mice were euthanized by carbon dioxide asphyxiation and weighed followed by exsanguination by terminal cardiac puncture. The blood placed into a 0.5 ml Greiner Bio-1 minicollection tubes (VWR, International L. L. C.) with ethylene diamine tetra-acetic acid (EDTA) for a complete blood count (CBC) by a commercial veterinary diagnostic laboratory (Antech Diagnostic Laboratory, New York, NY). The heart, lung (inflated with 10% neutral buffered formalin (NBF). All tissues except for the head, one rear leg, and sternum were immersion fixed in NBF. The sternum, head with the calvaria removed, rear leg with the skin removed were immersion-fixed and decalcified in Bouin’s solution (Ricca Chemical, Arlington, TX). All tissues were subsequently trimmed, placed in cassettes, processed to paraffin blocks, sectioned, and stained for hematoxylin and eosin by routine methods. The CBC results and slides were examined and photographed, and the figures made by Carmen J. Booth, D. V. M., Ph. D. Director, CPR Core, Yale University Department of Comparative Medicine.

### Lipid Nanoparticles (LNP)-crRNA packaging

crRNAs targeting Ttr were designed using the CRISPick website from Broad Institute^32^ and synthesized by IDT (Alt-R™) with IDT-proprietary RNA modification. crRNA was reconstituted in 300μl of RNase-free water and then 1.4ml of 25mM sodium acetate at pH 5.2 was added. Lipid mixture composed of 46.3% ALC-0315, 1.6% ALC-0159, 9.4% DSPC, and 42.7% Cholesterol and was diluted 4 times with ethanol before used. The LNP-crRNA was assembled using NanoAssemblr® Ignite™ instrument (Precision Nanosystem) following manufacturers’ instructions. The formulated LNP-crRNA was buffered exchanged to PBS using 30kDa Amicon filter unit and 40% (v/v) sucrose was added to the final concentration of 8% (v/v) as cryoprotectant. For quality control, the LNP-crRNA particle size was determined by DLS device (DynaPro NanoStar, Wyatt, WDPN-06) and the encapsulation rate and crRNA concentration were measured by Quant-iT™ RiboGreen™ RNA Assay (Thermo Fisher).

### Liver TTR protein targeting using LNP-crRNA

After genotype verification by PCR, 15 constitutive enAsCas12a-HF1 mice (8 males, 7 females, 8-14 weeks old) were randomly assigned, stratified by sex, to 3 treatment groups: NTC1 (3 males and 2 females), Ttr-cr1 (2 males and 3 females), and Ttr-cr2 (3 males and 2 females). For each group, LNP-crRNA containing 0.034µg of RNA was injected intravenously into each mouse. Serum samples were collected before injection (day 0) and day 6, day 12, and day 20 post injection via retroorbital blood draw. ELISA was performed on all serum samples after day 20 to measure the serum TTR level using Mouse Prealbumin ELISA kit (Abcam).

### Design and Cloning of AAV-CRISPR Vectors

The AAV-CRISPR vectors were designed to contain Cre recombinase under control of an EFS promoter for the induction of enAsCas12a-HF1 expression when delivered to LSL-enAsCas12a-HF1 mice. They also contain a truncated human NGFR (tNGFR) for better sorting of the infected cells. An open crRNA array cassette (double SapI cutting site) was controlled by a human U6 promoter for crRNA array cloning. pAAV-hU6-DoubleSapI-EFS-Cre-truncatedNGFR was generated by PCR and subcloned from pAAV-hU6-sgbbSapI-hTBG-Fluc-P2A-Cre. Guides targeting each of the four tumor suppressor genes (*Trp53, Pten, Apc,* and *Rb1*) were designed using the CRISPick website from the Broad Institute^32^ and individually tested in the NIH3T3-enAsCas12a-HF1 cell line using a lentiviral vector, pLenti-hU6-AsDR-EF1α-mCherry-2A-Puro. Working guides were concatenated into crRNA array, which was named crTSG, and cloned into the AAV-CRISPR vector.

### AAV Production and Purification

AAV-vector plasmid and AAV-crTSG were used for AAV9 production and purification using Lenti-X 293 T cells (Takara Bio). Each set of Lenti-X 293 T cells (6 150mm-plates) were transiently transfected with the AAV-vector or AAV-crTSG1, AAV9 serotype plasmid and pDF6 using polyethyleneimine (PEI). Lenti-X 293 T cells were cultured to approximately 80% confluency before transfection. Before transfection, medium was changed to 13ml serum free DMEM. The transfection mixture for each plate was prepared by first mixing 6.2μg AAV-vector/AAV-crTSG1, 8.7μg AAV9 plasmid, and 10.4μg pDF6 plasmid with 434μl of Opti-MEM and then adding 130μl of PEI. Transfection mixture was then incubated at room temperature for approximately 20 mins to allow transfection complexes to form. The transfection complexes were then added dropwise to each plate and 7ml of complete DMEM (D10) was supplemented to each plate 8hrs after transfection. Cells were harvested by cell scrappers and collected in 50ml canonical tubes. 1/10 volume (3ml chloroform for 30ml cells in DPBS) of chloroform was added to each tube and the mixture was vigorously vortexed for 5 mins to lyse the cells. 7.6ml of 5M NaCl was added to each tube, which was then vortexed for 10 seconds. After 5 mins of centrifugation at 3000g, the aqueous phase was collected to a new tube, in which 9.4ml of 50% (w/v) PEG8000 was added to precipitate virus particles. After 1hr incubation on ice, the virus particle was spun down at 3000g for 30 mins. The pellet was then resuspended with 5ml of HEPES buffer (50mM, pH8.0), followed by 1μl/ml Super Nuclease (SinoBiological; supplemented by 1μl/ml 1M Mg^2+^) treatment at 37°C for 30 mins. Pure chloroform was added at a 1:1 ratio to volume in each tube, which was vortexed for 10s, followed by 3000g centrifugation for 5 mins. The aqueous phase was collected, and the process was repeated until the aqueous phase became visibly clear. The final aqueous phase was isolated and passed through a 100-kDa Millipore filter unit. DPBS was added to washed and resuspend to a final volume of 0.5 ml. Genome copy number (GC) of AAV was titrated by real-time quantitative PCR using a custom Taqman probe targeting Cre.

### Tumor induction mediated by AAV intravenous (i.v.) and intratracheal (i.t.) injection

For i.v. injection, 12 LSL-enAsCas12a-HF1 mice (6 Males and 6 Females, 12 weeks old) were genotypically verified by PCR and randomly assigned to 2 treatment groups: AAV-vector (3 males and 3 females) and AAV-crTSG (3 males and 3 females). For each mouse, approximately 1.4 × 10^12^ genome copies of AAV (within the 200µl i.v. injection limit) were injected. Mice were monitored weekly for tumor growth and palpable tumor was observed starting from 3 weeks post injection. All mice were euthanized within 7 weeks post injection. Tumor tissue and major organs were harvested for IVIS spectrum imaging, gDNA extraction using Monarch^®^ Genomic DNA Purification Kit (New England Biolabs), and histology sectioning.

For i.t. injection, 12 LSL-enAsCas12a-HF1 mice (7 Males and 5 Females, 8-12 weeks old) were genotypically verified by PCR and randomly assigned to 2 treatment groups: AAV-vector (3 males and 3 females) and AAV-crTSG (4 males and 2 females). For each mouse, roughly 3 × 10^!!^ genome copies of AAV (within the 50 µl i.t. injection limit) were injected following a well-established protocol^44^. Mice were euthanized 10 weeks post injection. Lung and other major organs were harvested for IVIS spectrum imaging, gDNA extraction using Monarch^®^ Genomic DNA Purification Kit (New England Biolabs), and histology sectioning.

### Histology

Harvested tumor and major organs were placed in histology cassettes and submerged in 10% formalin for 24 hours and then transferred to 70% ethanol for storage. The samples were then sent to the Yale Department of Pathology for hematoxylin and eosin (H&E) staining and immunohistochemistry (IHC) staining (GFP, Ki67, CD45LCA, CD3, CD11b, Ly6B, F4/80).

### Nextera library preparation and sequencing

For each sample in the T7E1 assay, PCR products were labeled, amplified, and assigned barcodes using the Nextera XT DNA Library Prep Kit from Illumina. Each sample’s library underwent individual quality control and measurement using the 4150 TapeStation System (Agilent). This was followed by combining the libraries and purifying them with the QIAquick PCR Purification Kit (Qiagen). A subsequent round of quality control and measurement was performed again using the 4150 TapeStation System. The libraries were then sent to the Yale Center for Genome Analysis for sequencing on Novaseq 6000.

### Next Generation Sequencing data processing and quantify gene modification percentage

FASTQ reads were quality controlled by running FastQC v0.11.9 ^45^ and contaminations by the Nextera transposase sequence at 3’ end of reads were trimmed using Cutadapt v3.2^46^ (-a CTGTCTCTTATACACATCT -A AGATGTGTATAAGAGACAG). Processed reads were aligned to the amplicon sequence and quantified for insertions, deletions (indel), and substitutions using CRISPResso2 v2.1.3 ^47^. Specifically, we retrieved amplicon sequences, which were 150 ∼ 250 bp (according to the length of reads) flanking crRNA target sites, from the mm10 genome. A 5-bp window, centered on predicted enAsCas12a-HF1 cutting sites, was used to quantify genetic modification for each crRNA in both vector control groups and experimental groups (-w 5 -wc −2 -- exclude_bp_from_left 30 --exclude_bp_from_right 30). Allele frequency plots were generated with CRISPResso2 (--annotate_wildtype_allele WT --plot_window_size 12). Percent-modification data from each sample were aggregated for analysis and visualization in R.

### Generation of retroviral guide delivery system

The retroviral vector, pRetro-hU6-AsDR-doubleBbsI-EFS-Cre-mScarlet was generated by Gibson Assembly. hU6-DR-doubleBsmBI was amplified from pLenti-hU6-doubleBsmBI-EF1α-mCherry-Puro. The fragment was then subcloned into digested pRetro-hU6-doubleBbsI-Cas9Scaffold-EFS-Cre-mScarlet. The double BsmBI cut site downstream of the hU6 promoter was replaced by a double BbsI site due to the presence of multiple BsmBI cut sites in the backbone plasmid. Different retroviral vector variations (containing different crRNAs) were generated via BbsI digestion of the vector followed by ligation with crRNA arrays.

### Retroviral production and concentration

To generate retrovirus, 40 million HEK293-FS cells were plated into each 150mm tissue culture plate the night before transfection. HEK293-FS cells were cultured using the complete DMEM medium (DMEM supplemented with 10% FBS and 1% Pen/Strep). On the day of transfection, complete medium was replaced by 13ml pure DMEM and cells were allowed to incubate at 37°C for 30 mins to 1 hr. For each plate, 16μg of transfer plasmid (pRetro-hU6-DR-doubleBbsI/crRNA-EFS-Cre-mScarlet) and 8μg of pCL-Eco packaging plasmid was added to 440ul of Opti-MEM. After thorough mixing, 72ul of PEI was added to the mixture and vortexed such that PEI:DNA ratio was 3:1. After incubation at room temperature for 10 mins, the transfection complex was added dropwise to each plate. Cells were incubated at 37°C for 8hrs to allow transfection to take place. 8ml of complete DMEM medium was supplemented to each plate 8 hrs post transfection. Retrovirus in the supernatant was collected 48 hrs post transfection, followed by centrifugation at 3000g for 10 mins at 4°C to remove debris. The virus-containing supernatant was then concentrated by adding autoclaved 40% PEG8000 (m/v) to a final concentration of 8% PEG8000. Then, the mixture was incubated at 4°C overnight. The retroviral particles were spun down at 3000g for 15-30 mins and resuspended with 1ml of fresh complete RPMI (RPMI supplemented with 10%FBS, 1%Pen/Strep, 1%NEAA, 1%sodium pyruvate, 2% HEPES, and 50μM beta-mercaptoethanol). Retrovirus was then stored at −80°C before use.

### BMDC isolation and culture

The femur and tibia were harvested from LSL-enAsCas12a-HF1 mice and temporarily stored in ice-cold 2% FBS (Sigma) in DPBS (Gibco). Bones were then sterilized using 70% ethanol for 3 mins. Ethanol was then removed by washing with 2% FBS 3 times. Both ends of the bones were cut open with sterilized surgical tools and the bone marrow was flushed out using insulin syringes. RBCs were lysed with ACK lysis buffer (Lonza) for 2 mins at room temperature, washed with 2% FBS, and resuspended in complete RPMI medium. Cells were then filtered through 40 μm strainer and resuspended with complete RPMI supplemented with 25ng/ml murine GM-CSF (Peprotech) to a final concentration of 2 million cells/ml and plated in 24-well plates.

### CD8 T cell isolation and culture

The spleen was harvested from LSL-enAsCas12a-HF1 mice and placed in ice-cold 2% FBS (Sigma) in DPBS (Gibco). Single cell suspension was prepared by meshing spleen against 100 μm strainer, followed by lysis of RBCs using ACK lysis buffer (Lonza). After being washed with 2% FBS, spleenocytes were filtered through 40 μm strainer and resuspended in MACS buffer (0.5% BSA + 2mM EDTA in DPBS). CD8 T cells were isolated using Mouse CD8 T Cell isolation kit (Miltenyi) and then resuspended with complete RPMI to a final concentration of 2 million cells/ml. CD8 T cells were cultured in 96-well round bottom plates, and then activated by pre-coated plate-bound anti-CD3 (1μg/ml; Biolegend) and soluble anti-CD28 ((1μg/ml; Biolegend). The culture medium was supplemented with murine IL-2 (2ng/ml; Peprotech) and IL-12p70 (2.5ng/ml; Peprotech). After 72 hrs of activation, CD8 T cells were transferred to a new 96-well round bottom plate to remove activation and continued to be cultured in complete RPMI supplemented with a higher concentration of IL-2 (10ng/ml; Peprotech).

### Spin Infection

Polybrene was added to a final concentration of 10μg/ml 18-24 hours after cell plating. Cells were preincubated for 30 mins at 37°C. Concentrated retrovirus stock was diluted 2 times with complete RPMI (supplemented with 10 μg/ml of polybrene; 10 ng/ml IL-2 for T cell and 25 ng/ml GM-CSF). Cells were spun down at 900g for 90 mins at 37°C, followed by another 30 mins incubation at 37°C in incubator. Viral medium was then completely replaced by fresh medium and cells were cultured for additional 5-6 days.

### Antibody and flow cytometry

CD8 T cells were directly stained with antibody cocktails suspended in MACS Buffer (0.5% BSA and 2mM EDTA in DPBS) for 15 to 30 mins on ice. BMDCs were first incubated with anti-CD16/32 antibodies to neutralize IgG Fc receptors for 30 mins on ice prior to any surface staining. For all flow staining, LIVE/DEAD Fixable Near-IR Dead Cell Stain (Invitrogen) was used to selectively filter out dead cells from analysis. Anti-CD8a BV421 (Biolegend) was used to defined CD8 T cells, while anti-CD11c PE/Cy7 or anti-CD11b BV421 was used as a BMDC lineage-defining surrogate. mScarlet^+^ defined the infected cells. Staining antibodies for the targeted surface proteins on BMDCs included anti-CD24 APC, anti-CD11c PE/Cy7, anti-MHCII APC, and anti-CD80 BV421. Staining antibodies for targeted proteins on CD8 T cells included anti-CD27 BV605 and anti-Thy1 APC. Infected cells were enriched for all samples by sorting on BD FACSAriaII cell sorter with 4 lasers (405nm, 488nm, 561nm, and 640nm) to >80% purity. Flowcytometry data was analyzed by FlowJo software 10.8.2.

### Immune Cells Genomic DNA Extraction

For every 50,000 cells, 20ul of QuickExtract (Epicentre) buffer was added, and the suspension was incubated at 65°C for 30mins, then at 95°C for 5 mins to denature nuclease. Samples were vortexed thoroughly before PCR.

### Sample size determination

Sample size was determined according to the lab’s prior work or from published studies of similar scope within the appropriate fields.

### Replication

Number of biological replicates (usually n >= 3) are indicated in the figure legends. Key findings (non-NGS) were replicated in at least two independent experiments. NGS experiments were performed with biological replicates as indicated in the manuscript.

### Randomization and blinding statements

Regular *in vitro* experiments were not randomized or blinded. High-throughput experiments and analyses were blinded by barcoded metadata.

### Standard statistical analysis

Standard statistical analyses were performed using regular statistical methods. GraphPad Prism, Excel and R were used for all analyses. Different levels of statistical significance were accessed based on specific *P*-values and type I error cutoffs (e.g., 0.05, 0.01, 0.001, 0.0001). Further details of statistical tests are provided in figure legends and/or supplemental information.

### Code availability

The code used for data analysis and the generation of figures related to this study are available from the corresponding author upon reasonable request.

### Data and resource availability

All data and analyses for this this study are included in this article and its supplementary information files. Processed data for NGS or omics data are provided in Supplemental Datasets. Raw sequencing data will be deposited to NIH Sequence Read Archive (SRA) or Gene Expression Omnibus (GEO). Various materials are available at commercial sources listed in the Key Resources Table (KRT). Knock-in mice, vectors, cell lines, other relevant information, or data unique to this study are available from the corresponding author upon reasonable request.

